# Proteome analysis provides insights into sex differences in *Holothuria Scabra*

**DOI:** 10.1101/2024.03.26.586852

**Authors:** Chuhang Cheng, FeiFei Wu, Yizhi Xu, Chunhua Ren, Ting Chen, Shella Li, Peihong Shen, Fajun Jiang

**Affiliations:** Guangxi Key Laboratory of Marine Environmental Science, Guangxi Academy of Marine Sciences, Guangxi Academy of Sciences, Nanning, China; College of Life Science and Technology of Guangxi University, Nanning, China; CAS Key Laboratory of Tropical Marine Bio-resources and Ecology (LMB) / Guangdong Provincial Key Laboratory of Applied Marine Biology (LAMB), South China Sea Institute of Oceanology, Chinese Academy of Sciences, Guangzhou, China; School of Marine Sciences, Sun Yat-sen University, Zhuhai, China; School of Biological Sciences, University of Edinburgh, Edinburgh, England; BASIS international school, guangzhou, China

**Keywords:** *Holothuria scabra*, proteome, gonads, DAPs, sex differences

## Abstract

Sex-determining mechanism is still ambiguous for sea cucumber *Holothuria scabra* which only manifests gonochorism in gonad. In this study, proteomic analysis was employed to delineate sex-related proteins and genes in gonads of *H. scabra*, subsequently validated through Quantitative real-time polymerase chain reaction (qRT-PCR). A total of 5,313 proteins were identified via proteome sequencing. Among these, 817 proteins exhibited expression in both the ovary and testis, with 445 proteins displaying up-regulation and 372 proteins showing down-regulation. Furthermore, 136 and 69 proteins were identified as ovary-specific and testis-specific Differentially Abundant Proteins (DAPs), respectively. For the validation of 75 DAP coding genes, 9 genes were considered to be reliable. Notably, 25 ovary-bias proteins enriched in ribosome pathway strongly indicated the crucial role of ribosome in ovary. And 5S/18S rRNA ratio in *H. Scabra* may have potencial to establish a nondestructive method to distinguish sexes unambiguously. This study serves to furnish novel evidence pertaining to sex differences in *H. scabra*.

## 1. Introduction

*Holothuria scabra* is one of the nocturnal benthic species that customarily fed on algae and plankton, and it is widely distributed in the tropical waters (1). Sea cucumbers play significant roles in maintaining the pH balance and alkalinity of the seawater, contributing to the health of coral reef ecosystem (2). They accelerate bioturbation by ingesting the organic matter in the sediment and dissolve carbonate during feeding, thereby promoting the periodic cycle of calcium carbonate (3). *H. scabra* plays a critical role in nutrient cycling, participating as sedimentary nutrients in the form of food chain (4, 5).

Sexual dimorphism is the defining characteristic of organisms in which male and female reproductive organs occur in different individuals (6). The sex dimorphism phenotype is thought to be the result of differential gene expression profiles between genders, most prominently in gonads and germ cells. The mechanisms of gender determination and differentiation vary significantly across different metazoans due to repeated, independent lineage-specific evolution and rapid modification of potential molecular pathways (7). This variation is related to endocrine, neural, environmental, social, and ecological factors, including temperature, season, nutrition, and metabolic substances (8, 9). To expound upon the molecular mechanisms regulating sexual dimorphism, it is essential to examine the expression patterns of all sex-specific genes, particularly those involved in sexual-biased tissues (10).

In vertebrates, reproduction is controlled by the brain-pituitary-gonadal (BPG) axis (11). The growth, differentiation and maturation of follicular cells are induced by gonadal-expressed steroids and hormones (12, 13). Study on Half-smooth tongue sole (*Cynoglossus semilaevis*) has identified several ovary-related genes, such as ZPC, survivin, aquaporin, CPEB, O15, RacGAPO5, and O18. Some of these genes exhibit sexual dimorphism in the kidney, spleen, and muscle (14). Similar findings were observed in mice, where 27 genes displayed sexual dimorphism in the kidney (15). As for the invertebrate, numerous sex-related genes have been identified from a genome-wide scale for the sex differentiation mechanism in insects. Cyclin-related genes and serine/threonine-protein kinases (TSSKs) were suggested to be involved in spermatogenesis, while sex lethal (Sxl) and transformer-2 (tra) were proven to be associated with sex determination. The roles of ecdysone biosynthesis- and chorion-related genes in oogenesis have been elucidated (16-19). For Crustacea, differentially expressed genes, such as vitellin, vasa-like and gonadotropin-releasing hormone-like in *Litopenaeus vannamei* (20-22), cyclin A, cyclin B, transformer-2 (Tra-2) and cell division cycle 2 (Cdc2) in *Penaeus monodon*were (23-25), double-sex and mab-3 related transcription factor (Dmrt) in *Eriocheir sinensis*(26), proliferating cell nuclear antigen (PCNA) and heat shock protein 90 (Hsp90) in *Marsupenaeus japonicas* (27, 28), activated protein kinase C1 (RACK1) and cell apoptosis susceptibility (FcCAS) in *Fenneropenaeus chinensis* (29, 30) were found to be involved in sex determination and differentiation. In shellfish, sex differentiation is affected by double-sex-, SoxE and mab-3-related transcription factor (DMRT), β-catenin, forkhead box L2 (foxl2), and foxl2os (31).

Nevertheless, the lack of a high-quality genome severely hinders studies examining the sex determination mechanism of sea cucumber. Proteomics presents a promising alternative for the discovery of candidate proteins that exhibit significant differences between genders. Quantitative proteomics has been applied to aquatic animals to analyze crucial proteins and pathways involved in oogenesis and sex reversal (32, 33). Through label-free quantitativeproteomics (Labelfree) technology, this study examined the sex differences between the ovary and testis of *H. scabra*. The goal is to contribute to a comprehensive understanding of the mechanisms underlying sex determination and to acquire essential data on reproductive processes in *H. scabra*.

## 2. Materials and Methods

### 2.1. Samples collection

Wild adult *H. Scabra* specimens used in the experiment were sampled in Xuwen, Zhanjiang, Guangdong province, China (N20°42′, E109°94′). The gonads of the adult sea cucumbers were promptly dissected and categorized into two groups. One third of the specimen were preserved in Bouin fluid for sex conformation, the remaining were the frozen in liquid nitrogen and stored at -80°C for further sequencing.

### 2.2. Histological examination

The gonads were fixed in Bouin’s solution for 24 hours, gradually dehydrated using gradient ethanol, clarified with xylene and embedded in paraffin, paraffin-embedded tissue was sectioned into approximate 0.5cm^3^ cubes and then cut into 5-μm slices by a LEICARM2235 Slice Machine (Leica, Germany). The samples were stained with haematoxylin/eosin (H/E) and sealed with resin. Microscopic observations were conducted on sliced tissues using a Motic BA410 microscope (Leica, Germany).

### 2.3. Total protein extraction and digestion, Liquid Chromatography-Mass Spectrometry/ Mass Spectrometry (LC-MS/MS)

The tissue sample was pulverized under low temperature and mixed with protein lysis buffer. The resulting solution underwent a series of processes including ultrasonic lysis, DDT red and IAM reaction, acetone precipitation, resuspension, rinsing, and drying. Following this, protein dissolution buffer was added for dissolution, and protein quality tests were conducted using the Bradford Protein Assay Kit.

A 120 μg portion was taken from each protein sample mixed with Protein dissolution buffer TEAB buffer under 37°C, followed by enzymatic cutting and an overnight incubation. The solution was then treated with methanoic acid, centrifuged, and the supernatant was filtered using a C18 desalination column. Rinsing was performed three times with 0.1% methanoic acid and 4% acetonitrile, followed by elution twice with 0.1% methanoic acid and 4% acetonitrile. The eluates were merged, lyophilized, and subjected to LC-MS/MS analysis using the Q Exactive TM HF-X mass spectrometer. Spectrum was searched using PD2.2, Thermo. Inferential statistical analysis was carried out using Mann-Whitney Test for the results of protein assay, and the protein (p< 0.05) with a significant difference in experiment groups and control group is defined as differentially expressed protein (DEP). Program Interproscan-5 was used for gene ontology (GO) and InterPro (IPR) analysis of the Non-Redundant Protein Sequence Database (including SMART, ProDOM, ProSiteProfiles, Pfam, Panther and PRINTS). At the same time, the co-ortholog group (COG) and KEGG database were used to analyse the protein family and correspondent pathways. Through STRING-db server (http://string.embl.de/), possible protein-protein interaction is also predicted. Pathway enrichment analysis of GO, KEGG and IPR is then carried out.

### 2.4. Verification of sex differential genes

Total RNA of the whole transcriptome was extracted following the instructions provided in the TRIzol™ Reagent (Invitrogen) manual. The process involved tissue homogenization, chloroform extraction of RNA, isopropyl alcohol precipitation, washing with 75% ethanol twice, RNA dissolution in RNase-free ddH2O after precipitation, and measuring RNA concentration and purity using an Ultramicro spectrophotometer (Nanodrop 2000). Gel electrophoresis was then carried out for the detection of completeness of RNA samples.

For verification, 25 *H. scabra* specimens were collected from Dingda Seedling Farm, Wenchang, Hainan province, China (N19°45′, E110°78′) in July 2020. The gonads of these adult sea cucumbers were dissected, and sex determination was performed using the routine wax section method. The RNA extracted from the gonads underwent reverse transcription after concentration adjustment with the PrimeScript™ RT reagent Kit with gDNA Eraser (Perfect Real Time) (Takara, Japan).

From the proteome, 9 sex-specific genes were selected (primer is shown in table 3). Following the instructions from SYBP Premix Ex TaqTM II, a 25 μL Fluorescent-Quantitation PCR reaction system was employed (12.5 μL of SYBP Premix Ex Taq (2×), 1 μL of upstream primer, 1 μL of downstream primer, 2 μL cDNA and 8.5 μL RNase-free ddH2O). The cDNA templates originated from 4 female and 4 male *H. scabra*. Thermal Cycler Dice Real-Time System III was used for RT-qPCR with a two-step process. The reaction program included initial denaturation (95°C, 1 min), 40 cycles of 95°C for 5 seconds, and 60°C for 30 seconds, followed by signal collection under 72°C. There were 4 biological replicates and 3 technical replicates each for the genes and β-actin. The relative gene expression was normalized to β-actin by the comparative CT method. Pearson’s r correlation coefficient was calculated to evaluate the correlation between the qRT-PCR and proteomic analysis data.

## 3. Results

### 3.1. Histological Structure of the mature gonad in *H. scabra*

Total 30 *H. scabra* individuals were collected from Xuwen County, Zhanjiang City in September 2019 (Fig 1). HE staining was performed to characterize the structure of testis and ovary in mature males and females, respectively. In mature males, the genital atrium was filled with motile sperms which developed from spermatocyte (Fig 2a). Mature ovaries exhibited visible oocytes, forming an irregular polygonal shape due to the cells squeezing each other (Fig 2b). The total protein were extracted from six male and six female *H. scabra* individuals. These extractions were then sent to Novogene™ for LC-MS/MS detection analysis (Fig 2c).

**Fig 1.**
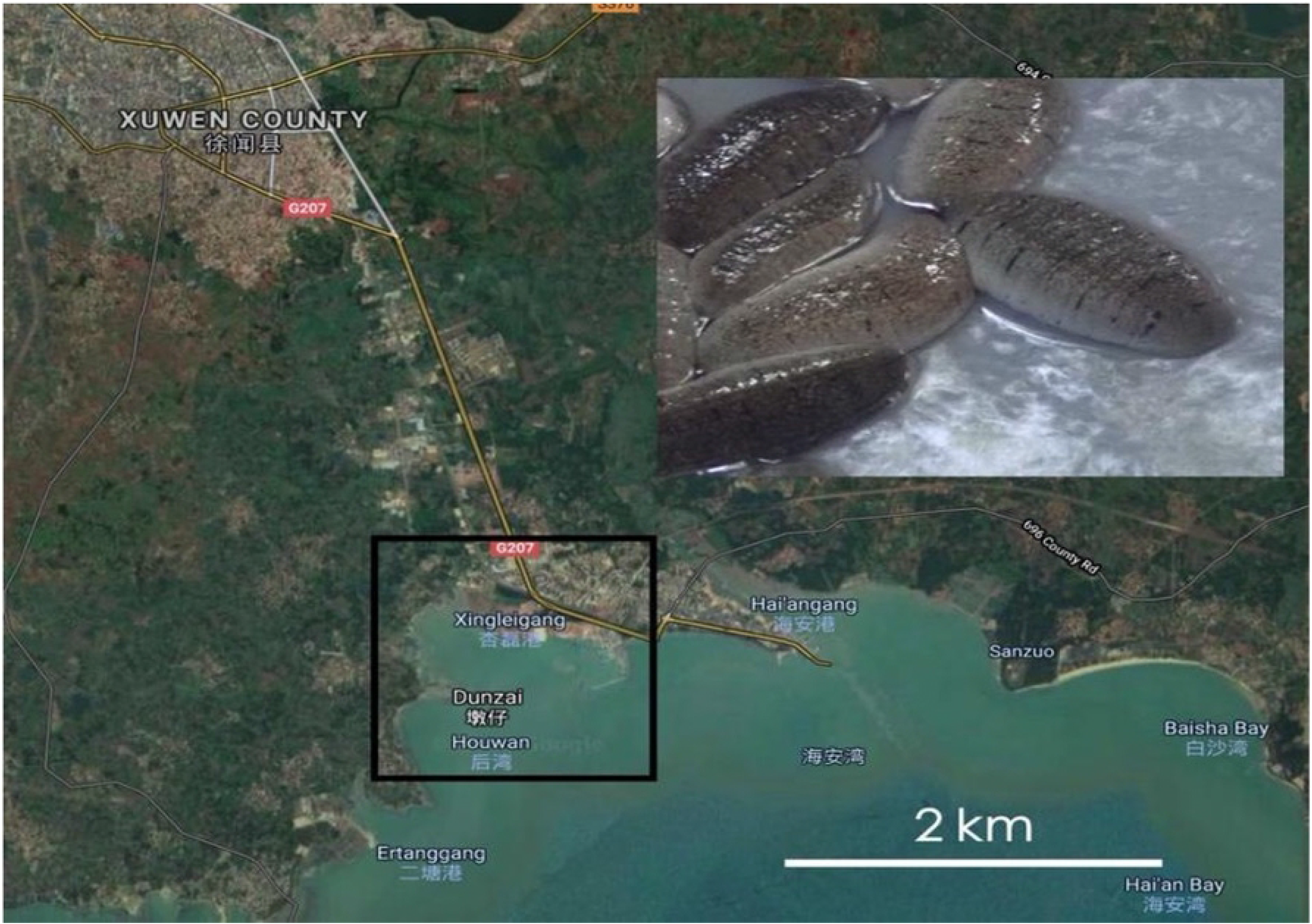
Location map of capture site for *H. scabra*

**Fig 2.**
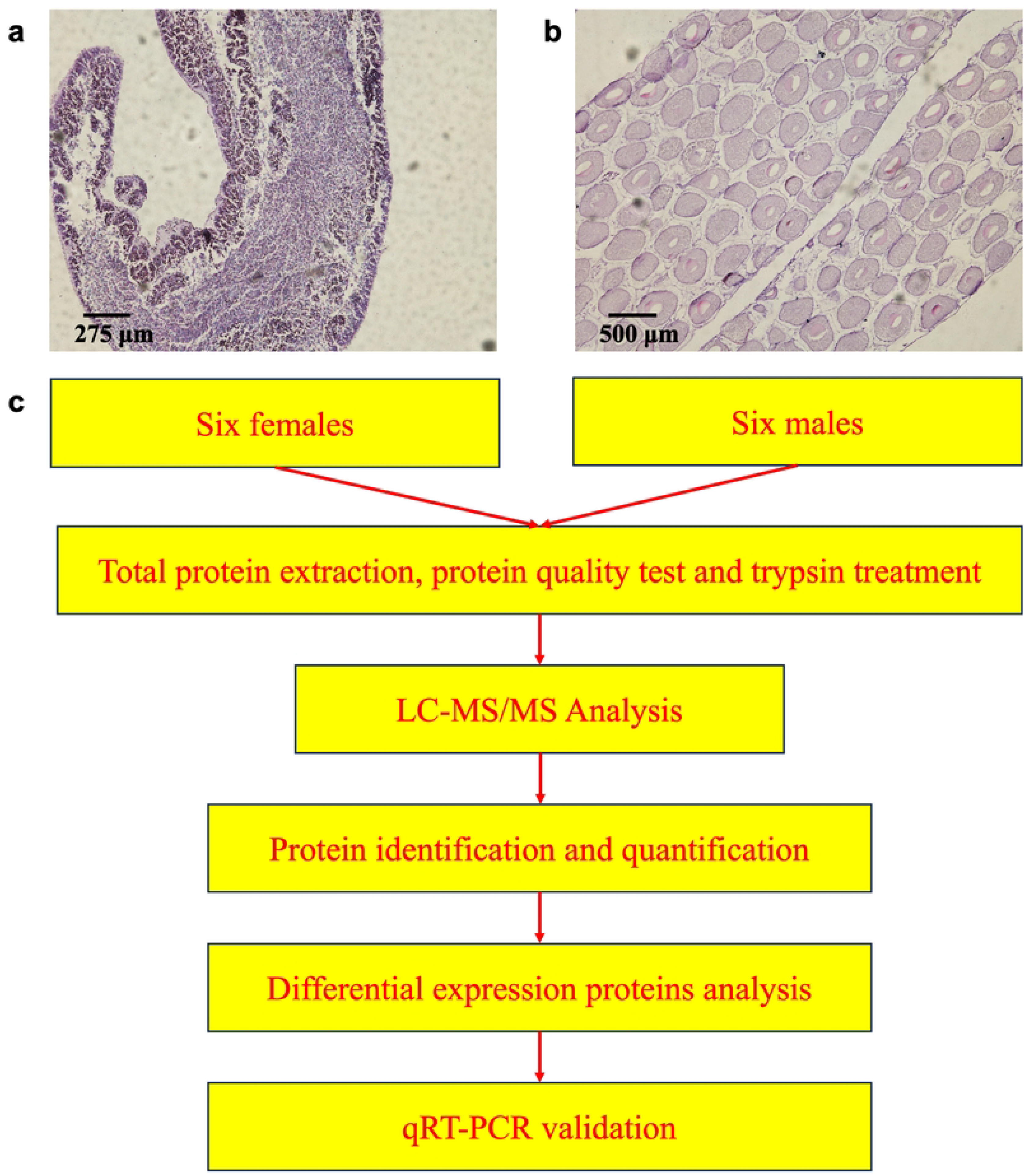
Gonad section of *H. scabra* and workflow (**a**) Testis (b) Ovary (**c**) Diagram of workflow for comparative proteomics between two sexes of *H. scabra*

### 3.2. Proteomic analysis data

#### 3.2.1. Statistics of Proteomic analysis data

Proteome sequencing yielded a total of 49,357 unique peptides, resulting in the identification of 5,313 proteins. The distribution of peptide length, protein coverage, and protein mass (S1 Fig a,b and c) demonstrated the accuracy and high reliability of the identification results. The results of Principal coordinates analysis (PCA) showed the significant separation between the proteins of two sexes of *H. scabra* (S1 Fig d).

#### 3.2.2 Functional annotation of all proteins

All the quantified proteins were functionally annotated using GO, KEGG, COG, InterPro (IPR), and subcellular localization (Fig 3). The Venn diagram shows a total of 4400 proteins annotated, with proximately 91.1% of them annotated by more than two databases. The GO enrichment analysis demonstrated that most proteins were enriched in molecular function, especially in the terms of protein and ATP bindings (S2 Fig). Furthermore, the KEGG pathway annotation showed that proteins identified in the gonads of *H. scabra* were mainly involved in metabolism, including carbohydrate, amino acid, lipid, nucleotide, and energy metabolism as presented in S3 Fig. The COG analysis classified the proteins into 26 functional categories, especially in general function prediction, translation, ribosomal structure, biogenesis, posttranslational modification, protein turnover, chaperones, and signal transduction mechanisms (Fig 3b). IPR annotation analysis mainly identified protein kinase domain, RNA recognition motif domain, and WD40 repeat-containing proteins. Subcellular location analyses were performed that cytoplasmic proteins (24.38%) and nucleus proteins (21.40%) comprised the largest proportion among the total proteins (Fig 3d).

**Fig 3.**
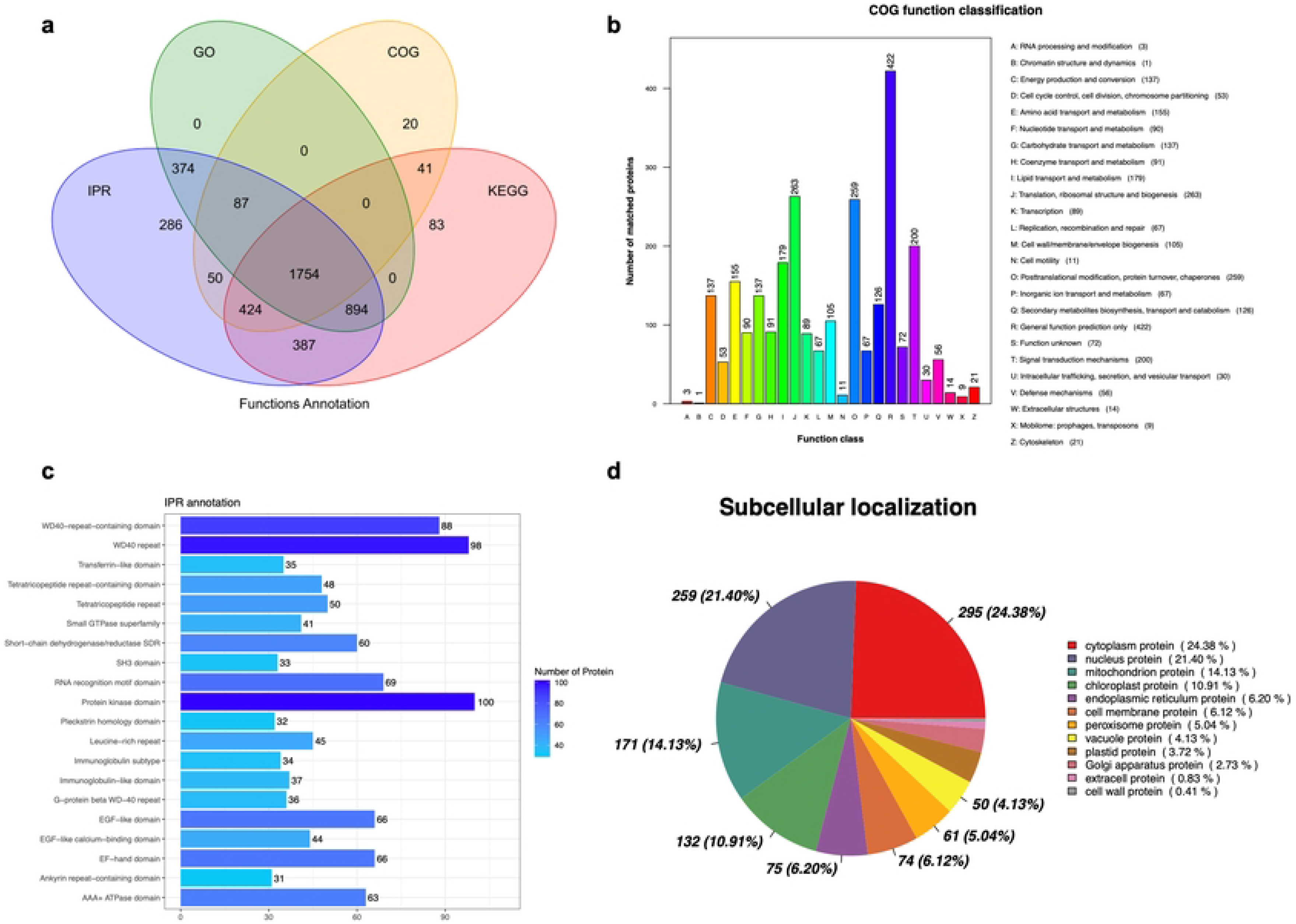
Functional annotation analysis (**a**) Wayne analysis of annotated proteins using different databases. (**b**) COG functional classification of all matched proteins. (**c**) IPR annotausing different analysis of of all samples. (**d**) The subcellular localization of all samples. GO gene ontology, COG: Cluster Cluster of of Orthologous Groups, IPR: InterPro, KEGG: Kyoto Encyclopedia of Genes and Genomes.

#### 3.2.3 Analysis of the DAPs associated with GO and KEGG Pathways

|log2(fold change) | >3 and p-value < 0.05 were set as a threshold to identify Differentially Abundant Proteins (DAPs). A total of 817 DAPs including 136 ovary-specific proteins and 69 testicle-specific proteins were obtained in samples after comparative analyses. Compared to the testis, there were 445 upregulated DAPs and 372 downregulated DAPs in the ovary (Fig 4a). Fig 4b shows significant difference between female and male gonads and the consistency of DAPs among every sample, which demonstrated the reliability of the data. Furthermore, the top 11 up-, and downregulated DAPs coding genes between two sexes were shown in Table 2 (p<0.001). And Table 3 listed 25 genes associated with sex and gametogenesis including egg coat matrix protein, egg bindin receptor protein 1 precursor, sperm flagellar protein and other sperm-associated proteins which were participated in the structure of gametes.

**Fig 4.**
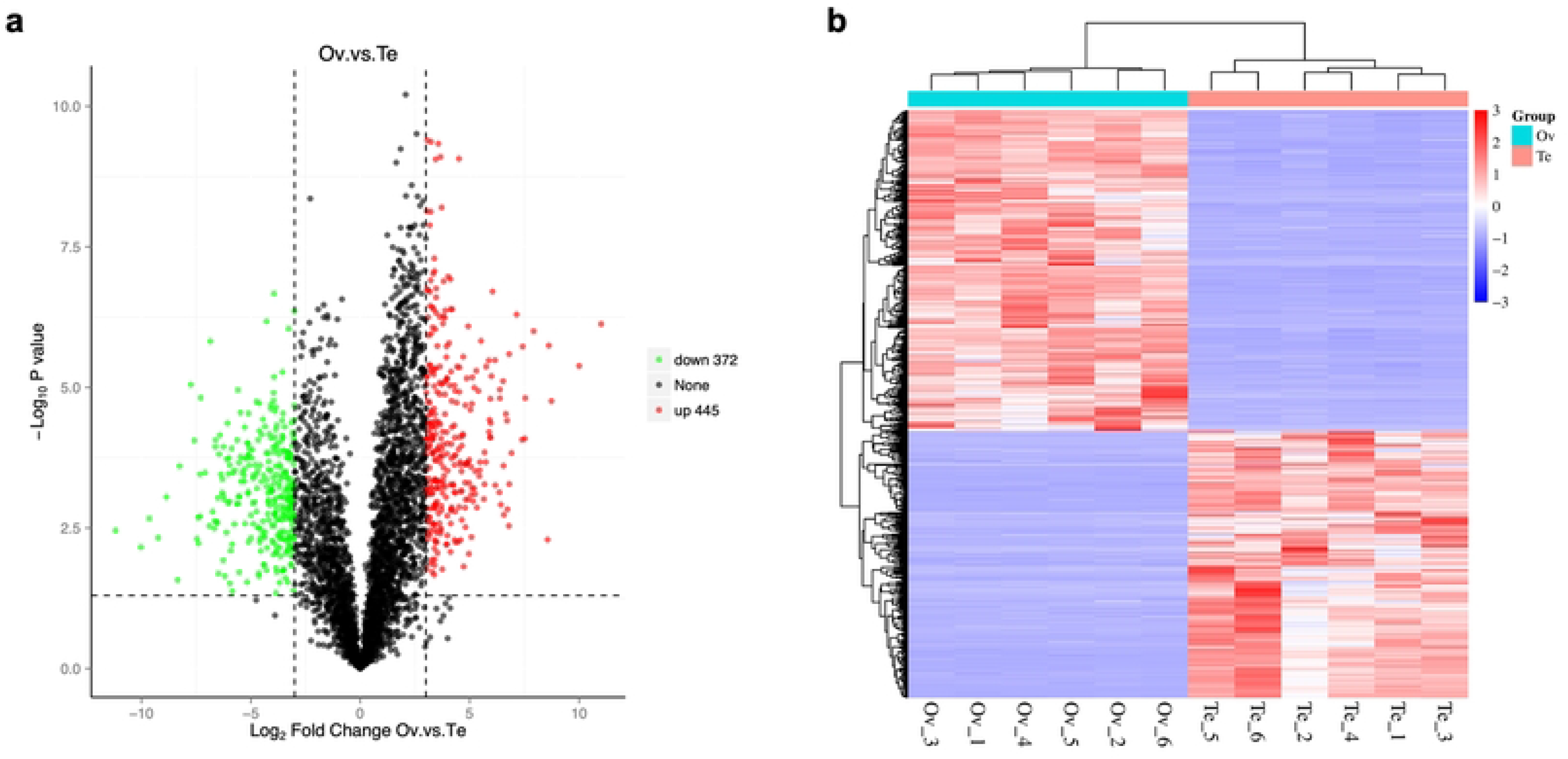
Differential expression analysis of proteins between ovary and testis **(a)** Volcano plot (**b**) Heatmap

Using gene ontology (GO), all DAPs were matched to 403 GO terms. The GO annotation chart showed the 29 enriched GO terms which categorized into three functional groups (S4 Fig). Fig 5 shows the top 10 ovary-bias and the top 10 testis-bias functional terms (P<0.05). Among them, Structural constituent of ribosome and Ribosome enriched 21 and 21 female-specific proteins respectively, which indicated strong ribosome-related activities in ovary. The GO terms related to membrane, membrane parts, and integral components of the membrane were enriched in many proteins upregulated in females. Most of upregulated proteins in testis were enriched under GO terms related to microtubule movement like microtubule-based process and microtubule-based movement. That is because the movement of sperms depends on the activities of flagella and the mature testicles are full of sperms. Ubiquinol-cytochrome-c reductase activity and ATPase activity, which involves in energy metabolism, were enriched many testis upregulated proteins.

**Fig 5.**
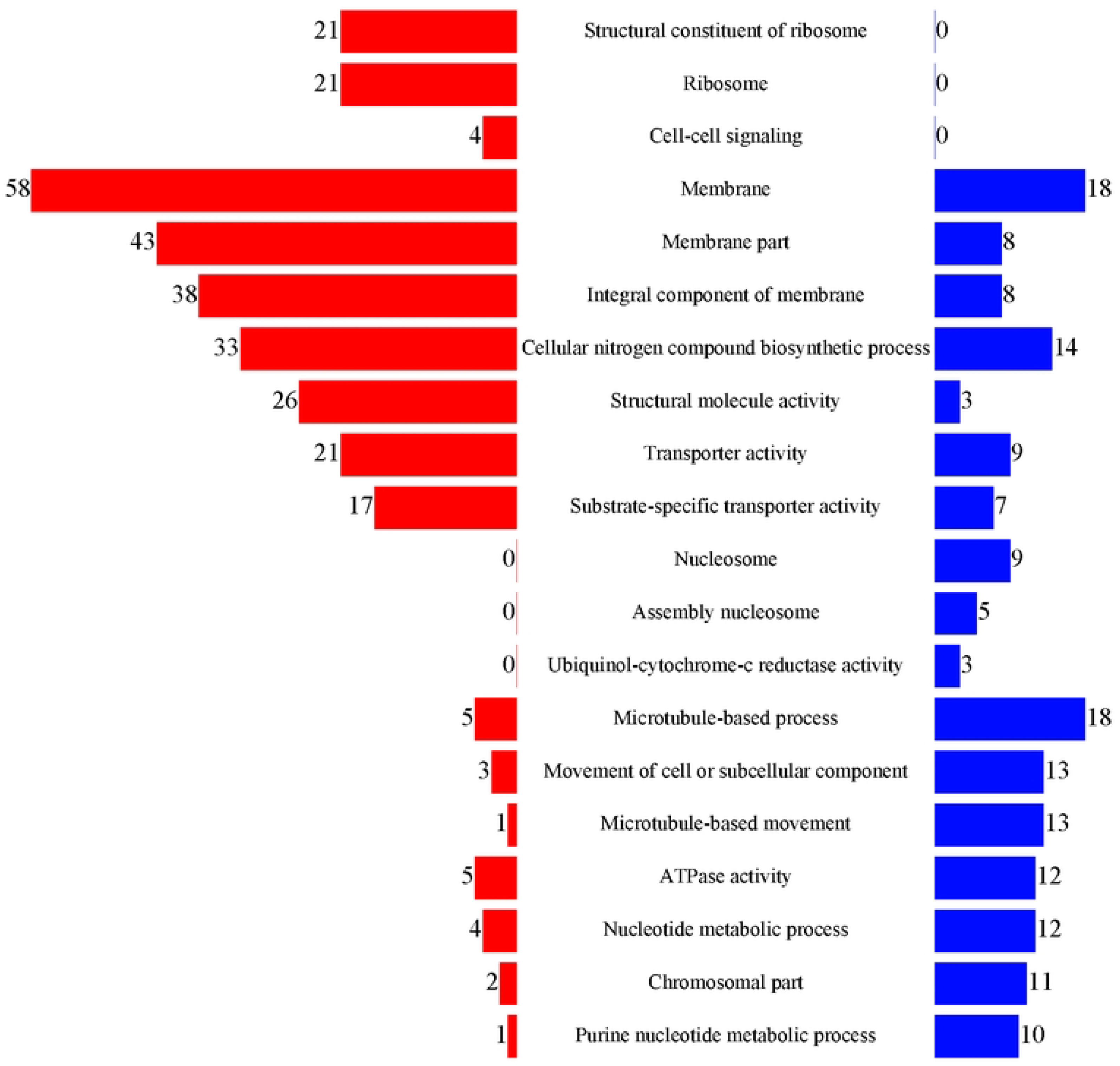
Number of DAPs in the 20 enriched GO terms (P<0.05). Red and blue color represents up-regulated and down-regulated proteins, respectively (Ov vs Te).

The KEGG analysis showed that differential proteins enriched the pathways associated with biochemical metabolic and signal transduction including sphingolipid metabolism, ABC transporters, phosphatidylinositol signaling system, amino sugar and nucleotide sugar metabolism, thiamine metabolism, beta-Alaine metabolism, and ribosome (Fig 6a). Notably, the ribosomal pathway enriched 24 up-regulated proteins (Fig 6b).

**Fig 6.**
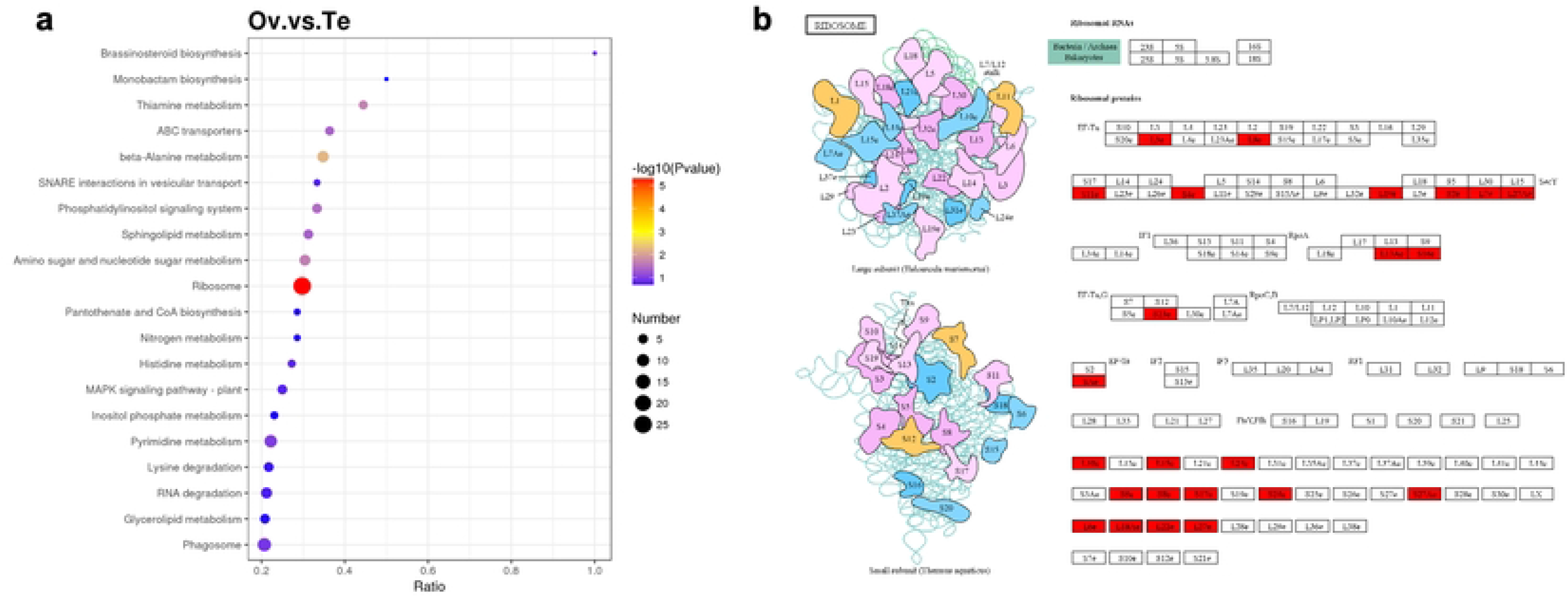
KEGG enrichment analysis (**a**) The differentially abundant proteins between ovary and testis. (**b**) Ribosomal pathway diagram. Red color represents up-regulated proteins.

### 3.3. Validation of Gene Expression from proteome by qRT-PCR

Totally, 75 selected proteins from proteome were to analyze by RT-qPCR according to the GO, KEGG annotation and Ov/Te expression pattern (S1 Table). As the results shown in Fig 7a and Table 4, seven of the genes in ovaries were verified significantly upregulated, and 2 of the genes were significantly downregulated. A strong correlation of qRT-PCR and proteomic analysis data was shown (Fig 7b), indicating the reliability of label-free quantitativeproteomics analysis to investigate the protein expression profiles of sex difference in *H. scabra*.

**Fig 7.**
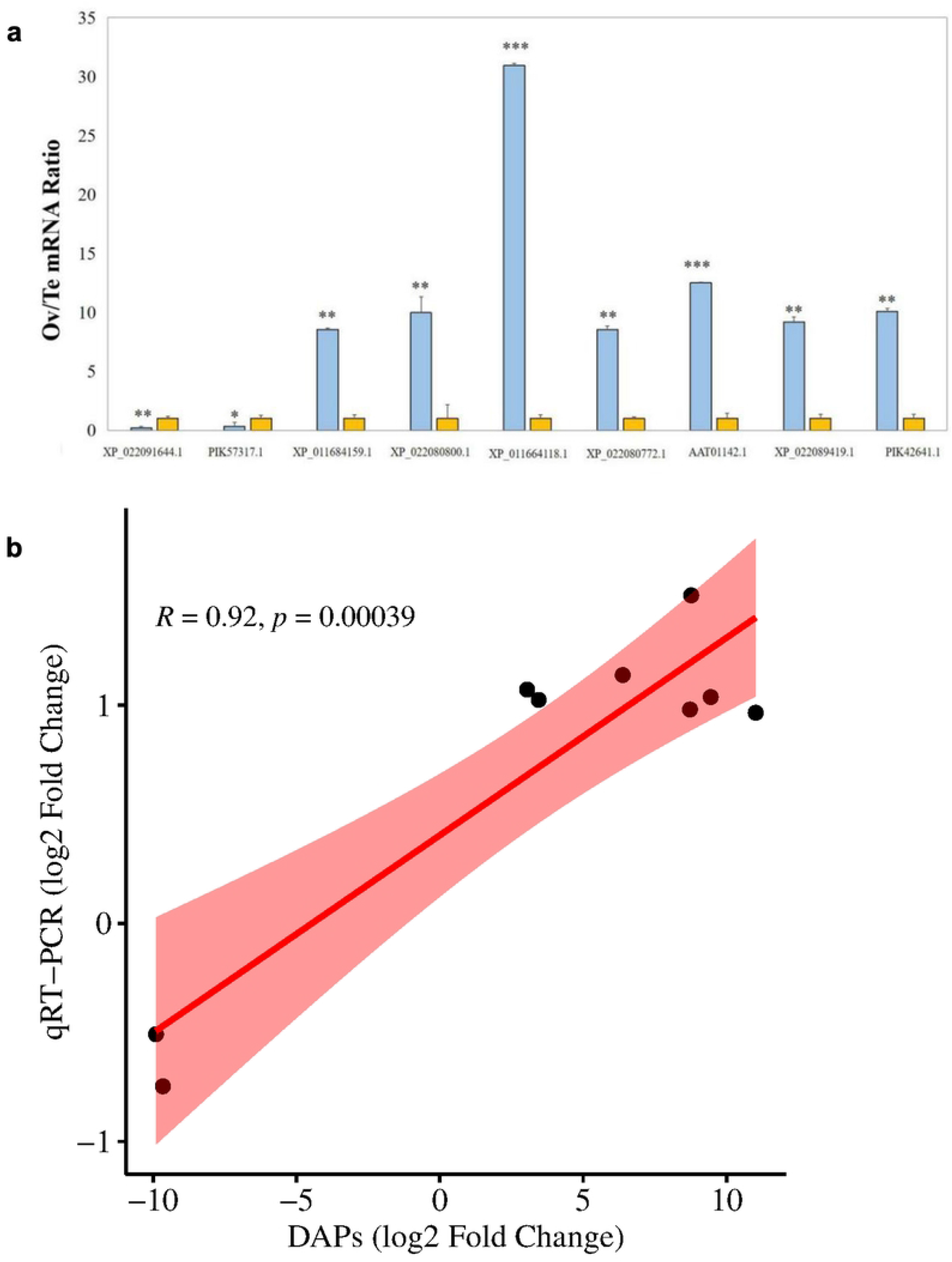
Quantitative real-time PCR (qRT-PCR) results for 9 coding genes (**a**) Relative expression levels of genes encoding DAPs. Y-axis denotes the fold change in gene expression of Ov/Te and all expression levels in \testis are set to 1. Blue represents Ovary, yellow represents Testis. (b) Pearson’s correlation analysis of qRT-PCR and proteomic data for DAPs.

## 4. Discussion

The sea cucumber *Holothuria scabra* is an economically important species of echinoderm in Asian market because of its high nutritional, pharmaceutical and economic value (34). The aquaculture of *H. scabra* is also popular for meeting the increased demands of market consumption and natural stock restoration (35). During sea cucumber culturing, usage of clear sexes parents will be benefit for the process of breeding and reproduction. However, the sexes of *H. scabra* cannot distinguish from appearance which may hinder the aquaculture of this species. And the sex determination of holothurians is still ambiguous. Previously, we have investigated the metabolomics profiles and sex makers of two sexes(36, 37). In this study, to enrich our knowledge of sex difference of *H. scabra*, the comparative proteomics between ovary and testis was performed using label-free quantitative method.

From the proteome analysis, 75 differential expression genes are identified in gonads of *H. scabra*. Verification through RT-qPCR was carried out, and the result indicates that there were 9 genes with significant differences. Upregulated proteins in male’s gonad, cilia- and flagella-associated protein, and sperm-associated antigen 8, play roles in spermatogenesis including sperm motility and microtubule formation(38, 39). Proteoliaisin is a protein that participates the assembly process of fertilization envelope in sea urchin(40). In echinoderms, proteoliaisin interacts with another protein ovoperoxidase to form a 1:1 complex. This complex inserts into the fertilization envelope to mediates hardening of the assembled envelope (41). In our result, ovoperoxidase and proteoliaisin were found significantly upregulated in ovary (showed in Table 1 and Table 3). That indicated the key components of formation of the fertilization envelope mainly exist in ovary and would interact after fertilization. Interestingly, hyalin, a large glycoprotein in the hyaline layer, was also detected high expression in female’s gonad (Fig 7a and Table 4). The hyaline layer locates underneath the fertilization envelope in zygote and play a role in blocking against polyspermy (42). Hyalin is also involved in regulating adhesive relationships as a specific cell adhesion molecule in the developing sea urchin embryo (43). Laminin subunit alpha, upregulated in ovary, can assemble into various laminin isoforms and is crucial for protein correct localization in the development of *Caenorhabditis elegans* (44). Those evidence suggested that the eggs carry many important proteins as preparation for early embryonic development in *H. scabra*.

**Table 1.**
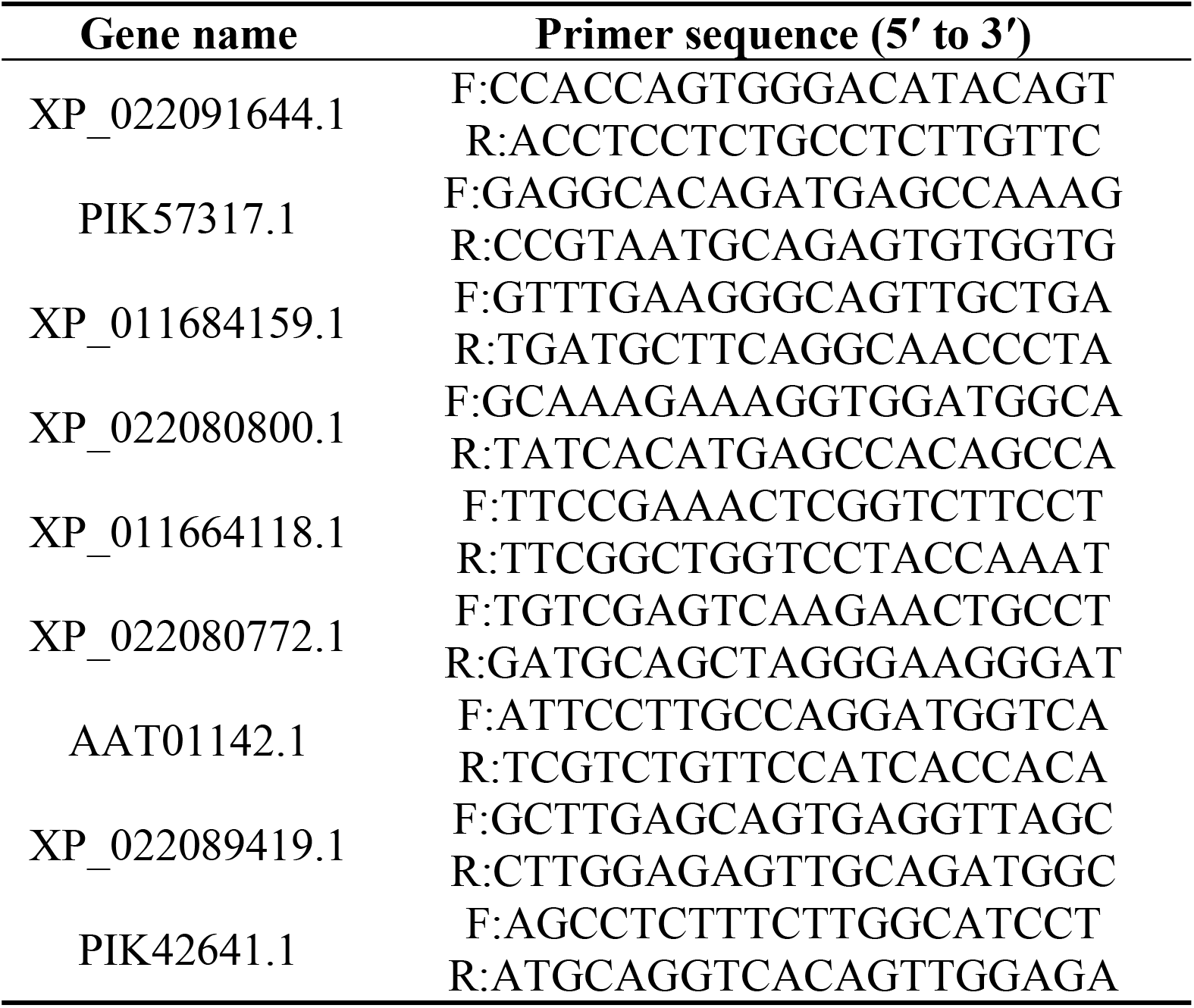
RT-qPCR primers for genes with significant difference from Proteome.

**Table 2.** 22 of the corresponding genes for significantly difference proteins between ovaries and testis.

| Gene | Description | log2FC | Trend | Pvalue |
| --- | --- | --- | --- | --- |
| PIK49186.1 | hypothetical protein BSL78_13967 | 10.00328 | up | 4.15E-06 |
| BAJ41227.1 | egg coat matrix protein | 7.924463 | up | 9.99E-07 |
| XP_013073410.1 | EGF and pentraxin domain-containing protein 1-like | 7.540273 | up | 1.55E-05 |
| AAB92242.1 | ovoperoxidase | 7.52235 | up | 8.14E-05 |
| XP_003726318.1 | GDP-fucose protein O-fucosyltransferase 1 isoform X1 | 7.144135 | up | 5.05E-07 |
| PIK39035.1 | hypothetical protein BSL78_24131, partial | 6.809735 | up | 5.25E-04 |
| PIK39115.1 | C-type lectin domain family 7 member A-like | 6.804186 | up | 2.55E-06 |
| PIK48450.1 | heme-binding protein 2 | 6.547302 | up | 2.48E-04 |
| PIK49839.1 | folate receptor gamma | 6.381332 | up | 1.56E-05 |
| AAT01142.1 | proteoliasin, partial | 6.379633 | up | 1.09E-05 |
| XP_011675803.1 | ATP-dependent RNA helicase cgh-1 | 6.044833 | up | 1.97E-07 |
| PIK49192.1 | enkurin | -6.10117 | down | 1.89E-04 |
| XP_022107427.1 | radial spoke head 1 homolog | -6.24921 | down | 1.14E-04 |
| PIK58530.1 | EF-hand domain-containing family member C2 | -6.29442 | down | 4.28E-04 |
| PIK56687.1 | egg permeability glycoprotein | -6.30547 | down | 7.44E-04 |
| XP_011667426.1 | tektin-4 isoform X3 | -6.36452 | down | 3.53E-04 |
| PIK38778.1 | radial spoke head protein 4-like A-like | -6.41209 | down | 7.07E-05 |
| PIK55121.1 | radial spoke head protein 9-like | -6.85095 | down | 1.51E-06 |
| XP_022081732.1 | creatine kinase U-type, mitochondrial-like | -7.08851 | down | 3.30E-04 |
| **PIK61874.1** | nucleoside diphosphate kinase-like 5 | -7.29379 | down | 1.53E-05 |
| **PIK57258.1** | creatine kinase, flagellar isoform X1 | -7.32068 | down | 3.50E-04 |
| **PIK40833.1** | ADP, ATP carrier protein | -8.85668 | down | 8.89E-04 |

**Table 3.** The list of sex and gametogenesis related DAPs between ovaries and testis.

| **Gene** | **Description** | **log2FC** | **Trend** | **Pvalue** |
| --- | --- | --- | --- | --- |
| **BAJ41225.1** | egg coat matrix protein | 8.616619037 | up | 1.80E-06 |
| **BAJ41227.1** | egg coat matrix protein | 7.924463339 | up | 9.99E-07 |
| **BAJ41226.1** | egg coat matrix protein | 7.423074104 | up | 1.88E-06 |
| **AAP44488.1** | egg bindin receptor protein 1 precursor | 3.301188988 | up | 3.76E-07 |
| **PIK35444.1** | sperm flagellar protein 1-like | -3.022187941 | down | 5.28E-04 |
| **XP_022090776.1** | cilia- and flagella-associated protein 44-like isoform X3 | -3.266191646 | down | 1.44E-04 |
| **XP_022105499.1** | cilia- and flagella-associated protein 57-like | -3.362091033 | down | 1.72E-03 |
| **PIK57014.1** | primary ciliary dyskinesia protein 1-like | -3.456240939 | down | 1.35E-03 |
| **PIK42503.1** | cilia- and flagella-associated protein 70 | -3.641234272 | down | 2.65E-04 |
| **XP_020604067.1** | cilia- and flagella-associated protein 20 | -3.892563548 | down | 2.65E-05 |
| **XP_022107058.1** | cilia- and flagella-associated protein 157-like | -3.912204831 | down | 1.80E-03 |
| **XP_022104631.1** | sperm-associated antigen 16 protein-like | -4.063316144 | down | 5.93E-04 |
| **PIK44114.1** | testis-expressed sequence 43 protein-like | -4.072539088 | down | 7.95E-05 |
| **PIK59902.1** | sperm-tail PG-rich repeat-containing protein 2 | -4.259111937 | down | 7.12E-03 |
| **XP_022086482.1** | cilia- and flagella-associated protein 58-like | -4.2825065 | down | 5.77E-04 |
| **PIK58729.1** | cilia- and flagella-associated protein 53 | -4.444913361 | down | 2.40E-03 |
| **PIK44983.1** | sperm-associated antigen 6 | 4.739561975 | down | 3.51E-04 |
| **XP_022102412.1** | dynein intermediate chain 3, ciliary | -4.947083538 | down | 1.80E-04 |
| **PIK34818.1** | testis-expressed sequence 36 protein-like isoform X1 | -5.533452907 | down | 1.94E-02 |
| **XP_022081058.1** | sperm-tail PG-rich repeat-containing protein 2-like isoform X1 | -5.845874193 | down | 4.22E-02 |
| **XP_022108113.1** | testis, prostate and placenta-expressed protein-like | -6.612017698 | down | 8.66E-05 |
| **PIK36385.1** | 63 kDa sperm flagellar membrane protein | -8.346589397 | down | 2.65E-02 |
| **XP_787834.2** | testis-specific serine..threonine-protein kinase 1-like | testis-specific | down | 0.00E+00 |
| **XP_022091644.1** | cilia- and flagella-associated protein 46-like isoform X7 | testis-specific | down | 0.00E+00 |
| **PIK57317.1** | sperm-associated antigen 8-like | testis-specific | down | 0.00E+00 |

**Table 4.**
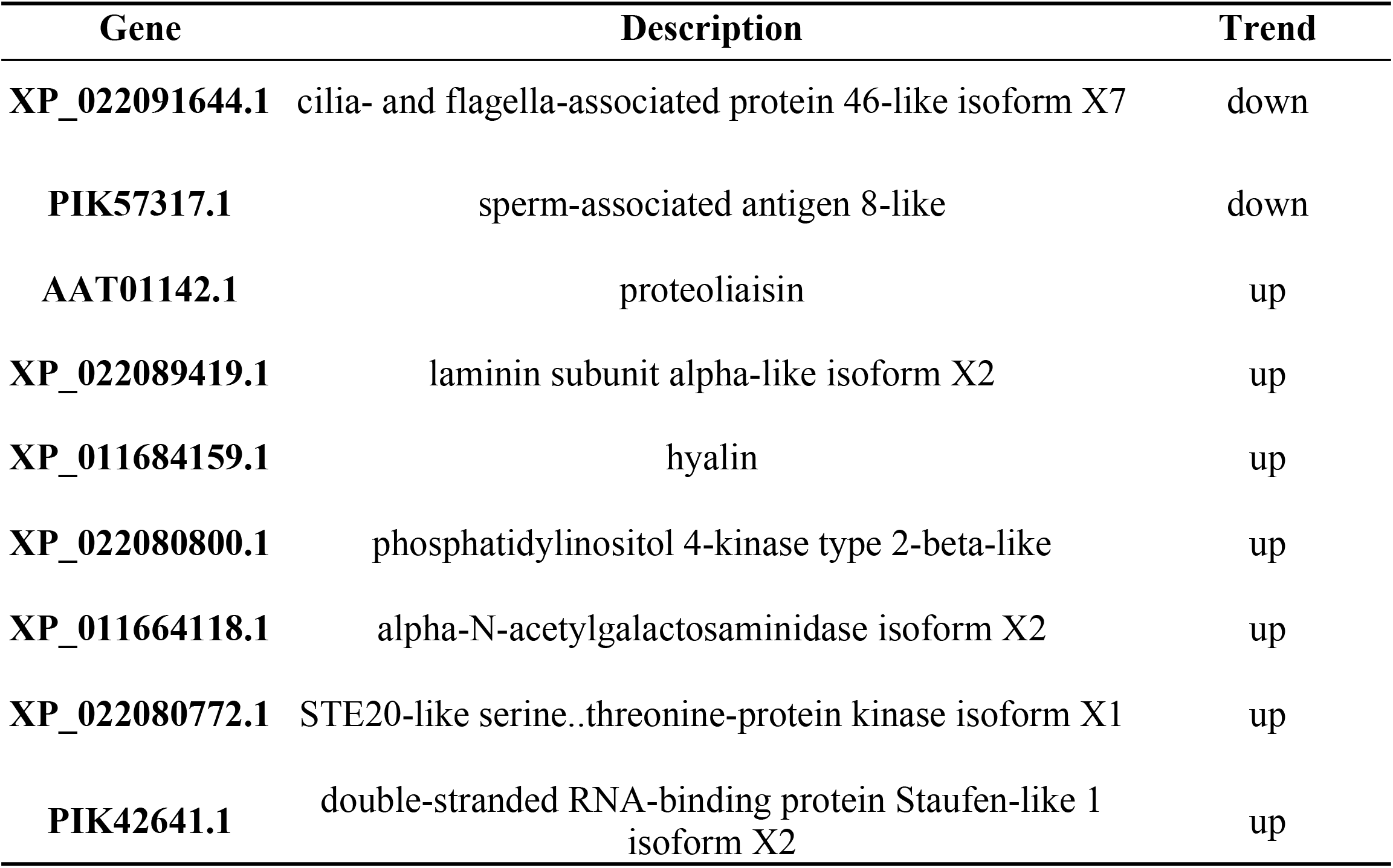
List of 9 sex differential genes after qRT-PCR verification.

In Go annotation, there were most ovary-specific proteins were associated function of structural constituent of ribosome and ribosome terms like 40S ribosomal proteins and 60S ribosomal proteins such as L3-like, L8, S6 and RPL15. Similar results were also found in KEGG pathway analysis. As showed in Fig 6b, the pathway of ribosome was enriched 25 upregulated proteins of ovary without any testis-bias proteins. Eukaryotes 80S ribosomes consist of a small (40S,including an 18S RNA and 33 proteins) and large (60S, including 25/28S, 5.8S, 5S rRNA and 49 proteins) subunit (45). Interestingly, many rRNA and ribosomal proteins have been linked with the ovarian development of aquatic animals. Previous studies in fish and reptiles have found the overwhelming accumulation of 5S rRNA in ovaries which indicated that its crucial role in oocytes (46-48). And 5S/18S rRNA ratio can serve as markers to distinguish sexes in fish (49). Ribosome protein S24 has been demonstrated as a potential stimulator in promoting the development of ovaries in east Asian river prawn *Macrobrachium nipponense* (50). Moreover, during oogenesis in the sea urchin *Paracentrotus lividus*, the expression of ribosomal protein S24 (RPS24) is increased (51). In our study of gonads proteome, we also found many significantly upregulated ribosome proteins in ovary which indicated that the ribosome plays a crucial role in ovary. The oocytes accumulate reserve substances for proper development of the embryo and ribosome contribute considerably to the synthesis of proteins in this process.

Most testis-specific proteins were classified in nucleosome and assembly nucleosome (Fig 5). In mammals, successful production of mature sperm involves the process of chromatin organization which make itself become highly compacted in the sperm head (52). Chromatin remodeling of the male genome during spermiogenesis relies on nucleosome. A nucleosome consists of a section of DNA that is wrapped around a core of histone proteins which is the basic repeating subunit of chromatin. During spermiogenesis, nucleosome transfer from a histone-based structure to a mostly protamine-based configuration which lead the chromosomes to become compact and condensed (53, 54). We have identified several histones like histone H1, H3, H4-like and H5 which were significantly high expression in testis (S1 Table). That result demonstrated those proteins is crucial to generating a viable male gamete in *H. scabra*. Motility and morphology are also thought to be indispensable for the fertilizing ability of sperm. Microtubule-based processes in spermatogenesis involve in sperm head shaping and sperm flagella development (55). Thus, it can be understood that sex differential proteins related to microtubules are enriched in testis (Fig 5).

The utilization of label-free quantitative proteomics allowed us to conduct a comparative proteomics analysis between two sexes of *H. Scabra* and to investigated protein candidates that might be involved in sex differences. In present study, we identified and verified 2 downregulated and 7 upregulated genes which involve in sperm motility and assembly process of fertilization envelope respectively. According to functional analysis, ribosomal proteins, membrane proteins, membrane part proteins and integral component of membrane were upregulated in ovary proteome while nucleosome, assembly nucleosome, microtubule movement, ubiquinol-cytochrome-c reductase activity and ATPase activity related proteins were high expression in testis. Notably, ribosome pathway only enriched 25 ovary-bias proteins which strongly indicated the crucial role of ribosome in ovary. And 5S/18S rRNA ratio in *H. Scabra* should be studied in the future to establish a nondestructive method to distinguish sexes unambiguously. Overall, our proteome results provide a novel insight for the study of sex mechanism in *H. Scabra*.

## Supporting information

**S1 Fig**. Statistics of the quantitative proteomics data of different samples. (a) The distribution of peptide lengths in all samples. (b) The distribution of peptide coverage in all samples. (c) Numbers of proteins with different masses in all samples. (d) Principal coordinates analysis of four types of individuals. PC: principal coordinate.

**S2 Fig**. GO enrichment analysis of labeled proteins

**S3 Fig**. KEGG enrichment analysis of labeled proteins

**S4 Fig**. GO enrichment analysis of abundant proteins in two comparison groups

**S1 Table**. Selected proteins list for qRT-PCR validation.

## Acknowledgments

Thanks to CAS Key Laboratory of Tropical Marine Bio-resources and Ecology (LMB) / Guangdong Provincial Key Laboratory of Applied Marine Biology (LAMB), South China Sea Institute of Oceanology for providing the experimental platform, reagent consumables, etc. We thank Shanghai NewCore Biotechnology Co., Ltd. (https://www.bioinformatics.com.cn, last accessed on 20 Feb 2024) for providing data analysis and visualization support.

